# Spatial Connectivity Pattern of The Human Brain’s Action Network

**DOI:** 10.1101/2025.10.12.681882

**Authors:** Xiu-Xia Xing, Xi-Nian Zuo

## Abstract

This study proposes a spatiotemporal connectome-based framework to characterize the human brain’s action network. Unlike temporal connectivity (TPC), this approach leverages full functional connectivity profiles derived from large-scale neural wave dynamics, termed spatial connectivity (SPC). We applied SPC to map the action network for the first time in a non-Western young adult cohort. Our results delineate the network’s detailed functional architecture across the cerebral cortex, cerebellum, and subcortical nuclei. Compared to TPC, SPC is more robust to global signal regression when characterizing anticorrelation between the action and default networks. Critically, this intrinsic antagonism reflects a fundamental energy-saving balance in the brain’s dynamical system. Moreover, SPC reveals that the salience/parietal memory network resides at the interface of this antagonism, exhibiting weak but highly variable connectivity, a position that enables rapid state switching between external action and internal contemplation. These findings offer a mechanistic view of the brain’s resting “dark energy” and suggest a blueprint for brain-inspired AI. Embedding a dual-system opponent architecture, balanced by a flexible switching hub, may foster adaptive, energy-efficient intelligence. All derived high-resolution SPC patterns and computational code are publicly shared to promote open science (https://ccndc.scidb.cn/en).

## Introduction

Quantifying functional connectivity between brain regions requires a multi-principal framework that reflects the biological and topological mechanisms of network formation^1-3^. Traditional approaches rely on temporal correlation between regional time series, thus measuring temporal connectivity (TPC)^4^. This method may overlook the complex spatial, developmental, and topological constraints that shaped neural wiring. Generative models show that network connectivity follows principles such as wiring economy (minimizing connection length) and homophily (preferential linking between nodes with similar connectivity profiles)^5,6^. Functional connectivity therefore emerges from a trade-off between spatial constraints and topological optimization. We should incorporate whole-brain spatial embedding to account for physical distance and connectivity homophily. Meanwhile, long-range, but weak connections play a critical role in integration^7-9^. Including the subcortical and cerebellar regions while preserving weak connectivity can improve connectomics reliability^10^. These considerations warrant a spatial connectivity method for quantifying functional connectivity with full-brain profiles.

Classic TPC treats relations between time series as statistical measures rather than physical objects. This leads to difficulties in interpretating weak relationships. Recent advances in cortical traveling waves offer new insights into the spatiotemporal patterns of neural activity reflected by resting-state fMRI (rfMRI) signals^11-13^. The rfMRI time series are observations of brain waves traveling along cortical surface. These waves are governed by principles of wave dynamics, including interference, superposition, and resonance. Large-scale functional networks can be modeled using neural field theories and coupled wave oscillators^14^. These models are based on the underlying connectional structure, particularly the exponential distance rule and long-range horizontal fibers, which serve as the substrate for wave propagation. This framework explains how synchronized oscillations emerge and travel across the cortex. It provides a mechanistic basis for the observed spatiotemporal profile of functional connectivity. From this viewpoint, the TPC map of a given seed area is a spatial profile. Each position’s TPC value quantifies the similarity between the seed’s rfMRI wave and the traveling wave at that position. Thus, TPC becomes a spatial snapshot of wave phases. The entire TPC spatial map has biological significance for coordinating activity across distributed regions. It facilitates efficient communication between functional systems, enabling integration of sensory input with internal states^15-18^. In summary, shifting from statistical magnitude to physical wave phases allows TPC metrics to be interpreted beyond magnitude. This enables full-space mapping to become a physical observation of traveling wave dynamics, providing a spatial basis for quantifying functional connectivity^19,20^.

In the present work, we propose a novel method of functional connectivity: spatial connectivity (SPC). Specifically, SPC employs the full-brain TPC map of an area without thresholding. This map preserves all connectivity and serves as a more valid characteristic of a functional system. It reflects activity propagation in both space and time, compared to time series observed only in the temporal domain. To test SPC, we investigated human brain dynamics by mapping the action-mode network (action network)^21^. This network functions oppositely and counterbalances the default-mode network (default network). The action network was originally defined using TPC and task activation, which fail to capture its spatiotemporal, system-level operations^22^. Traditional approaches based on statistical thresholds overlook the temporal evolution and propagating wave dynamics that likely underlie the action network’s role. To address these limitations, we investigated the SPC of the action network. We compared the network’s SPC and TPC with respect to their spatial maps and cortical network associations.

## Materials and Methods

### Open Datasets & Preprocessing

We used open datasets from the Chinese Human Connectome Project (CHCP)^23^. The acquisition of CHCP rfMRI data was consistent with the HCP protocol^24^. Prior reports demonstrated that 30 minutes of data provides reliable and optimal design for brain-wide studies^25^. CHCP Therefore obtained two 15-min rfMRI scans (AP and PA phase encodings) for all participants. All preprocessing steps were identical between CHCP and HCP with an updated version of the HCP pipeline (https://github.com/Washington-University/HCPpipelines/releases/tag/v4.0.0). After quality control, we included 217 healthy young adults in the subsequent analysis. Their preprocessed individual rfMRI data have been openly shared through the Science Data Bank (https://ccndc.scidb.cn/en), as described in the original publications^23^. We note that no global signal regression was performed during preprocessing. This step is known to introduce anticorrelation in traditional TPC analysis^26^.

### Group-level Temporal and Spatial Connectivity

We temporally trimmed each preprocessed individual dataset and applied and variance normalization. We then submitted the data to principal component analysis (PCA) at the group level (MIGP)^27^. Group PCA compressed the datasets using the weighted top 2500 spatial eigenvectors for further renormalization and reweight to form group-level timeseries (*\**.*dtseries*). According to rfMRI protocols, these PCA-compressed MIGP time series capture intrinsic spontaneous brain activity at the group level (i.e. 30 minutes).

We correlated the MIGP-timeseries to form the average dense TPC connectomes (*\**.*dconn*) at the group level (91282×91282 entries). Specifically, let *g*_*i*_ denote the *i*-th gray coordinate (*i* = 1,2, … ,91282), and *t*_*i*_ = [*ts*(*g*_*i*_, 1), *ts*(*g*_*i*_, 2), … , *ts*(*g*_*i*_, 2500)] denote standardized timeseries measured at the coordinate *g*_*i*_. The TPC connectome’s entry at (*g*_*i*_, *g*_*j*_) is defined by the cosine similarity between *t*_*i*_ and *t*_*j*_, as shown Equation (1).

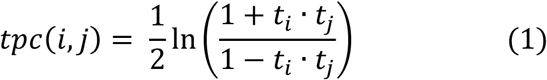

Here, *s*_*i*_ = [*tpc*(*i*, 1), *tpc*(*i*, 2), … , *tpc*(*i*, 91282)] denotes standardized full-brain TPC map seeded at coordinate *g*_*i*_. The spatial connectivity between *g*_*i*_ and *g*_*j*_ is defined by the cosine similarity between *s*_*i*_ and *s*_*j*_, as shown in Equation (2).

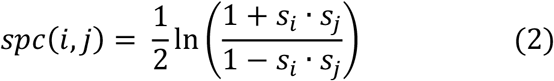

This yields the full-brain SPC connectome’s entry at (*g*_*i*_, *g*_*j*_) based on the average dense TPC connectome at the group level. Specifically, we treat the full TPC profile of a given spatial location as a snapshot of traveling wave dynamics measured by the spontaneous slow oscillations (SSO) from rfMRI scans. We then standardized cosine angles between each pair of snapshots to produce the dense SPC connectome at the group level.

### Network Connectivity Mapping

As illustrated in Figure 1, we selected four seeds of the action network for each hemisphere. We used the adjusted DU15NET parcellation in the **fsLR_32k** HCP grayordinate space (91,282 elements: 59,412 for cerebral cortex, 17,853 for cerebellum, and 14,017 for subcortical nuclei)^28^. In this parcellation, we replaced the original somatomotor A/B networks with the somatomotor and somato-cognitive action networks (SCAN)^29^. Table 1 summarizes the spatial and label information of the eight seeds.

**Table 1.** Information of the four seeds in the human action network.

| full name | dorsal Anterior Cingulate Cortex | anterior Insula Frontal Operculum | anterior Prefrontal Cortex | Supramarginal Gyrus |
| --- | --- | --- | --- | --- |
| abbreviation | dACC | aINFO | aPFC | SMG |
| XYZ (LH) | -8.6, 7.6, 37.4 | -32.4, 6.5, 11.1 | -29.6, 42.2, 25.3 | -54.2, -37, 30.7 |
| XYZ (RH) | 9.7, 6.9, 38.8 | 33.6, 8.5, 10.7 | 32.9, 41.6, 24.9 | 54.6, -28.5, 33.2 |

**Figure 1.**
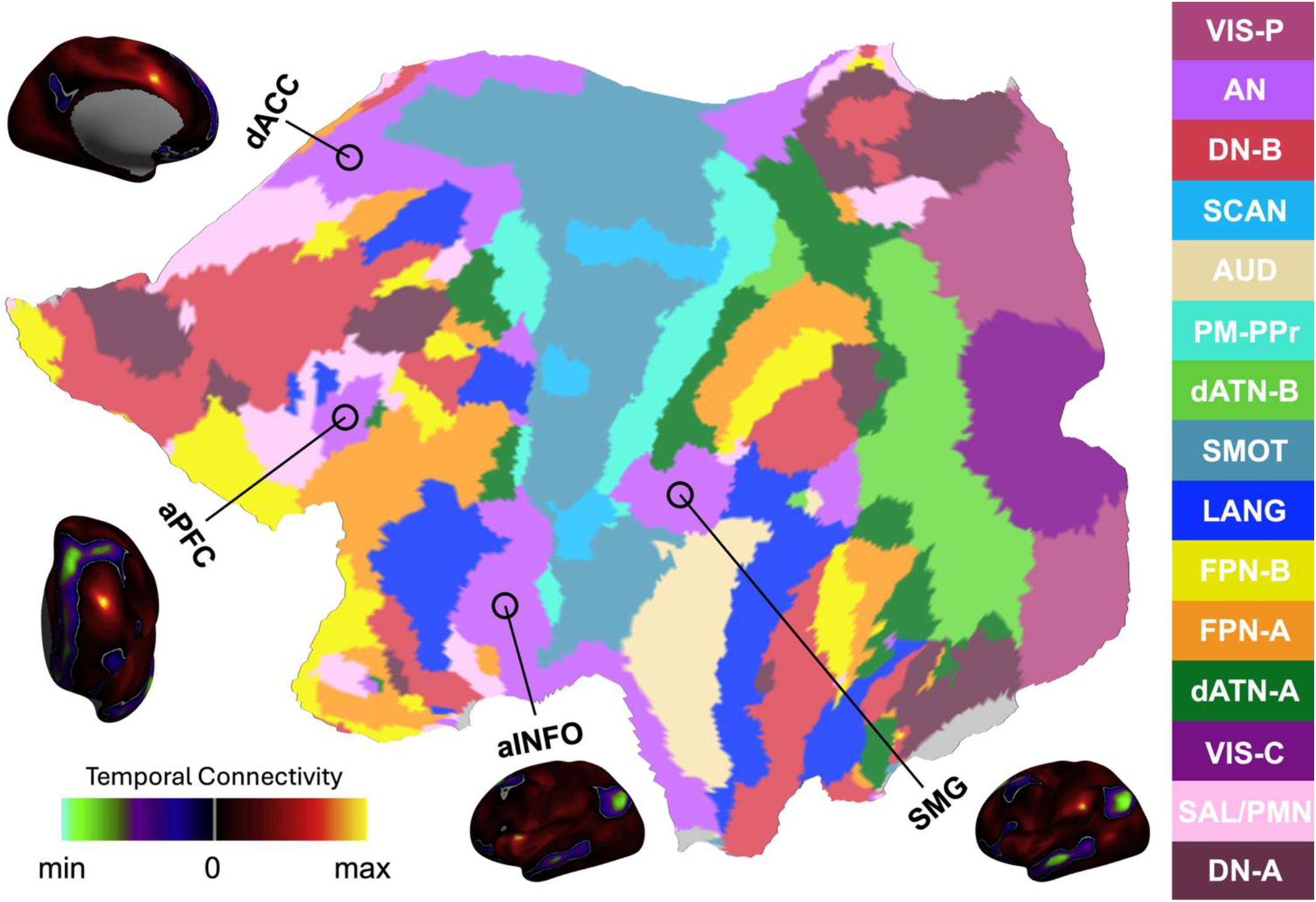
Methodology of action network connectivity in the human brain. Four seed regions are circled on the flat map of the left hemisphere: dorsal anterior cingulate cortex (dACC), anterior insula/ frontal operculum (aINFO), anterior prefrontal cortex (aPFC) and supramarginal gyrus (SMG). The original label of action network in DU15NET is cingulo-opercular (CG-OP) named by its anatomical structures network and has been updated by its functional correspondence ‘action network’ (AN). Meanwhile, the two somatomotor networks (SMOT-A and SMOT-B) are merged and then divided into the somato-cognitive action network (SCAN)^29^ and another remaining somatomotor network (SMOT). Other twelve networks are premotor-posterior parietal rostral (PM-PPr), salience/parietal memory network (SAL/PMN), dorsal attention-A (dATN-A), dorsal attention-B (dATN-B), frontoparietal network-A (FPN-A), frontoparietal network-B (FPN-B), default network-A (DN-A), default network-B (DN-B), language (LANG), visual central (VIS-C), visual peripheral (VIS-P), and auditory (AUD). The TPC maps of the four seed areas are rendered onto the fsLR_32k cortical surfaces and visualized, respectively.

For a given seed, we extracted its two‐step neighbors’ SPC maps from the group‐level dense SPC connectomes. We then calculated the median map of these SPC maps to quantify the seed network connectivity. We averaged all eight seeds’ network SPC maps to obtain the final action network SPC map. We derived the action network TPC map using the same approach for methodological comparison. Similarly, we used an identical procedure to derive the default network’s SPC and TPC maps^30^ as network‐level references (Figure S1 and Table S1).

## Results

We performed identical analyses on both CHCP and HCP datasets and observed highly similar results. Here, we presented the main results from CHCP dataset, with HCP-derived results as in the Supplementary Materials.

### Action Network Connectivity Maps

Figure 2 shows network connectivity maps of the human brain’s action network. Traditional TPC revealed only positive connectivity of the action network with the 15 cortical networks, with values varying across networks. Specifically, sensorimotor, visual and auditory networks exhibited the strongest AN-TPC. Default and control networks showed the weakest AN-TPC. Interestingly, the most variable AN-TPC existed between action network and salience/parietal memory network.

**Figure 2.**
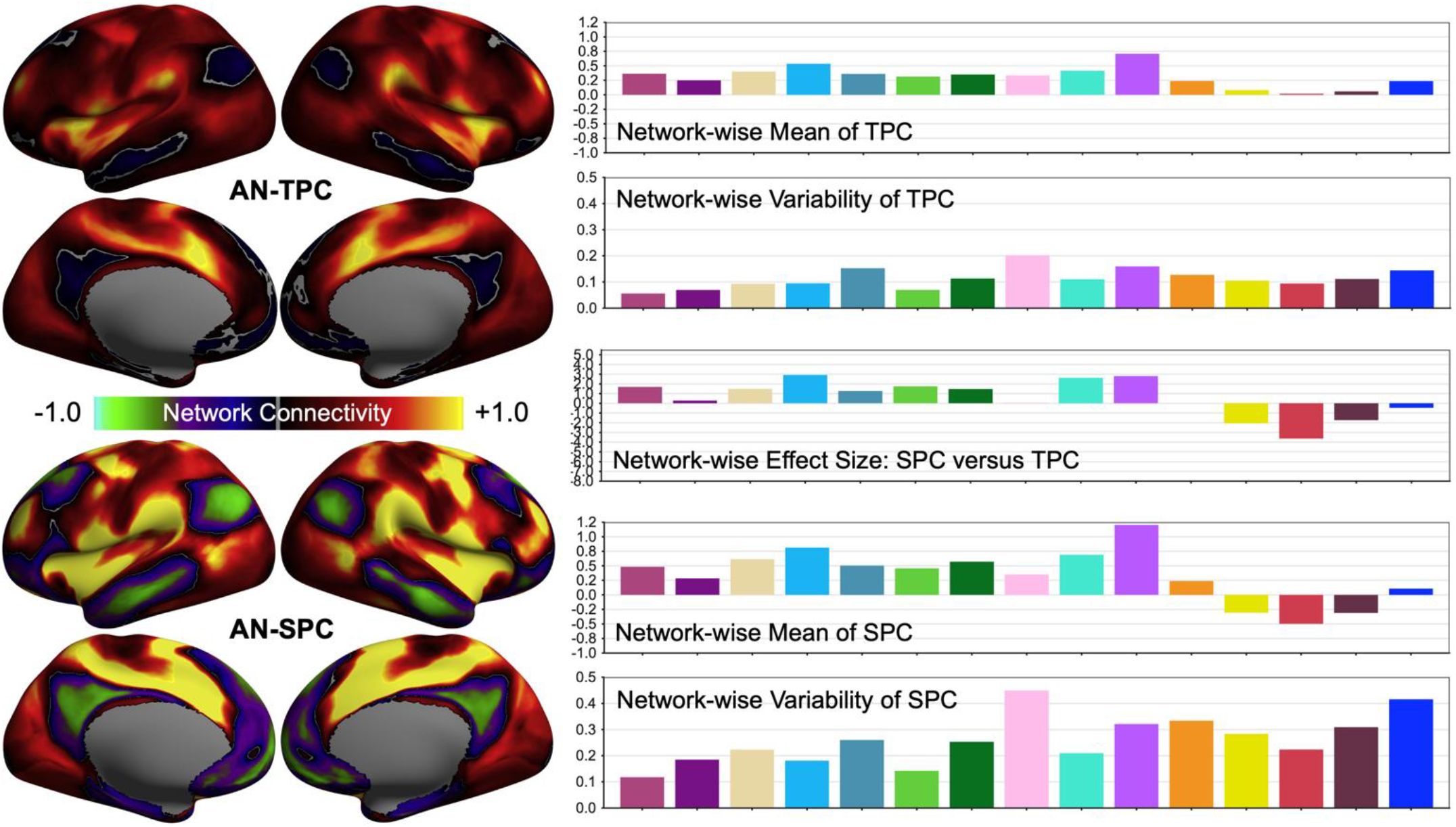
Mapping action network connectivity in the human brain. The network connectivity maps of action network (AN) are derived from the CHCP rfMRI data. Its temporal connectivity (AN-TPC) maps are rendered onto the standard cortical surfaces (top left). The network-wise mean or variability of TPC values are plotted as in the bar graph across the adjusted DU15NET networks (top right): the same color annotation used as in Figure 1. Same surface maps and bar plots are presented for spatial connectivity of action network (AN-SPC) in bottom left and bottom right. To compare maps between TPC and SPC, we plot the network-wise effect size of SPC minus TPC in the middle right panel.

These observations were reproducible for AN-SPC. Beyond enhancing positive connectivity, AN-SPC produced obvious negative values (i.e., anticorrelation) in default and control networks. We confirmed these differences through analysis of network-wise effect size on SPC-TPC, as described in Equation (3) and shown in Figure 2. Similar findings for default network are presented in Figure S2.

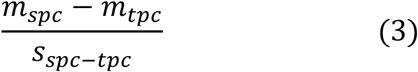

### Spatial Pattern of Action Network Connectivity

We visualized the upper 20% of the AN-SPC with positive or negative values (Figure 3). This represents synchronous or asynchronous functional communications of spatial-temporal dynamics in the cerebral cortex, cerebellum and subcortical nuclei. This method precisely delineates the cortical topology of the action network, outlining the boundaries of common networks (the violet color), namely the cingulo-opercular or salience/ventral network^23,28,31^. Negative connectivity of the action network was well distributed in default network areas, echoing their ‘Yin-Yang’ communicative dynamics. This profile of integration of the action- default integration was mirrored in the cerebellum and subcortical structures (mainly putamen and thalamus). The complete default network’s SPC map is shown in Figure S3 and demonstrates a highly similar but mirror pattern of network spatial connectivity.

**Figure 3.**
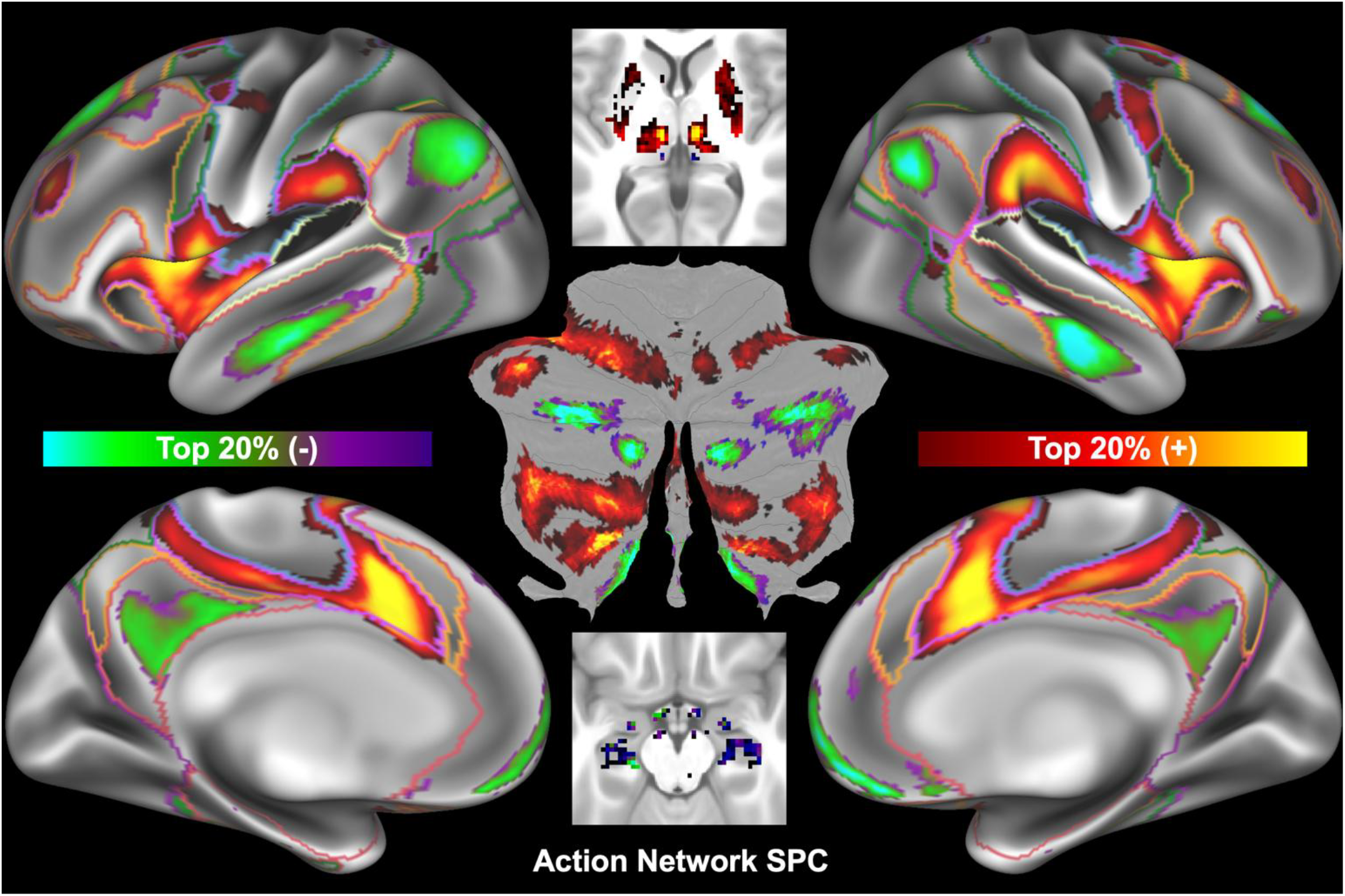
Spatial connectivity pattern of action network in human brain. The SPC map of action network was derived from the CHCP rfMRI data. Only the top 20% network SPC values were displayed for the three brain structures: cerebral cortex, subcortical nuclei and cerebellum. An integrative version of the DU15NET parcellation was used to outline boundaries of the seven large-scale networks derived by the CHCP team: visual (purple), somatomotor (blue), auditory (cream), dorsal attention (green), action (voliet), control (orange) and default (red). The action network SPC map was depicted in the two axial slices (z=4mm and z=-17mm) to demonstrate the high affinity of putamen and thalamus. This SPC map was also rendered on the flattened surface of the cerebellum.

## Discussion

The reconceptualization of the cingulo-opercular network as the action-mode network marks a significant advance in systems neuroscience^21,29,32^. This action network is proposed as the central circuit that enables the brain to enter an ‘action-mode’ - a functional state characterized by heightened arousal, external attention, goal-directed planning, motor execution, and responsiveness to action-relevant feedback such as pain and errors. It integrates executive control, motor planning, sympathetic drive, and viscerosensory processing under a unified framework centered on goal-directed behavior. A key feature is its strong anticorrelation with the default network^30^, positioning the two networks in a yin–yang relationship across the brain’s functional continuum^33,26^. Recent evidence from human and animal studies supports the action network’s role in motor plasticity, arousal regulation, action initiation, and integration with the SCAN^29^, which coordinates whole-body movement and physiology^32^.

### Population Diversity and Generalizability

While the action network framework is compelling, we must acknowledge the demographic and methodological constraints underlying its current definition. Nearly all foundational studies cited in support of the action network are based on Western neuroimaging datasets^21^, primarily from the United States and Europe. Our work demonstrates the action network’s anatomy and function in Chinese young adults for the first time. This compensates the canonical maps of action network derived from the ABCD study^34,21^, which limits the generalizability to adult populations. While action network has been shown to drive developing functional topography from childhood to adolescence in previous work^35-39^, our work fills this gap by generating and reproducing the action network maps in both CHCP and HCP (Supplementary Materials) adult samples. The absence of large-scale, cross-cultural, and lifespan-oriented functional atlases^3^ means that the action network’s stability, maturation, and cultural modulation remain poorly understood. Future research should prioritize replicating action network mapping in diverse adult cohorts and cultural contexts to validate its universality or identify population-specific variations^23,40-43^. The action network represents a major integrative model for understanding goal-directed behavior. Yet its current definition rests on geographically and developmentally narrow data. Expanding research to include global and adult samples is essential.

### The Action-Default Anticorrelation as a Dynamic Principle

The robust anticorrelation between the action and the default networks, consistently revealed by our SPC framework across both CHCP and HCP cohorts, is more than a statistical curiosity. We argue that this antagonistic relationship embodies a fundamental principle of the human brain as a complex dynamical system^44,45^. In such systems, opposing forces often coexist in a delicate balance. This push–pull interplay enables the system to operate near a critical state, where information transmission is maximized, metabolic cost is minimized, and rapid transitions between distinct functional modes (external action versus internal reflection) become computationally efficient^46^.

From an energetic perspective, this yin–yang equilibrium carries profound implications^47-50^. The brain consumes approximately 20% of the body’s resting energy budget despite accounting for only 2% of its mass. It must continuously allocate resources between competing functional demands. The action–default antagonism likely serves as a dynamic energy-saving mechanism. When one network is actively engaged, the other is relatively suppressed, reducing simultaneous high‐cost coactivation and preventing wasteful interference. This oscillatory metabolic partitioning may be a key evolutionary adaptation. It allows the brain to sustain its high baseline activity, the so‐called “brain dark energy”, while retaining rapid responsiveness to environmental challenges. Thus, the anticorrelation we observe is not merely a reflection of task‐negative versus task‐positive systems. It is a direct manifestation of how the brain’s intrinsic energy landscape shapes its functional architecture^51^.

### Implications for Brain Dark Energy

Our findings offer a fresh perspective on the elusive concept of brain dark energy, the persistent, intrinsically generated neural activity that consumes the majority of cerebral energy even without external stimuli. The action–default antagonism, captured by SPC without recourse to global signal regression, suggests that this resting‐state activity is not a homogeneous background hum. It is a structured, opponent‐process dynamic. The asynchronous alternation between the two networks may represent a low‐frequency “breathing” of the brain’s energy field, continually reconfiguring the system’s global state to prepare for unpredictable future demands. Thus, such dark energy is not chaotic noise. It is the spatiotemporal signature of a self‐organizing, antagonistic oscillator that maintains the brain in a poised state, ready to switch between introspective and extroverted modes with minimal energy expenditure.

This biological principle of antagonistic equilibrium holds salient lessons for modern artificial intelligence (AI). Most current deep learning and reinforcement learning systems lack any intrinsic opponent‐process architecture. They are trained to optimize a single objective, often without internal tension or self‐regulating trade‐offs. By contrast, the brain’s action–default antagonism provides a natural regularizer that prevents runaway excitation or pathological inertia, promoting adaptive flexibility, generalization, and robust performance under uncertainty. To align artificial systems more closely with human cognition^52-54^, we propose that future AI architectures should incorporate a built‐in “functional opposition” between two complementary subsystems. One would handle active goal‐directed processing. The other would manage off‐line consolidation, simulation, and imaginative planning (reminiscent of the “contemplative system” driven by dark energy). This dual‐system design, governed by a dynamic balance akin to the action–default antagonism, could enhance energy efficiency, facilitate continual learning without catastrophic forgetting, and endow agents with intrinsic motivation mirroring human curiosity and self‐reflection.

### Methodological Advance of SPC

The validity of this antagonistic relationship has been debated in conventional fMRI studies. Traditional TPC measures are notoriously sensitive to preprocessing choices, particularly global signal regression, which artificially introduces negative correlations. Thus, TPC renders the observed anticorrelation method‐dependent and potentially spurious. Our proposed SPC overcomes this limitation. By defining connectivity based on the full spatial profile of traveling wave snapshots rather than on point‐wise temporal correlations, SPC yields robust anticorrelations between the action and default networks without global signal regressing. This confirms that the action–default antagonism is not a preprocessing artefact but an intrinsic property of the brain’s spatiotemporal dynamics, a genuine feature of its self‐organized, wave‐propagating architecture^11-20^. SPC provides a more biologically faithful and preprocessing‐resilient tool for probing large‐scale network antagonisms.

### The Role of the Salience/Parietal Memory Network

Our results unveil an intriguing finding regarding the salience/parietal memory network (SAL/PMN)^56-60^. Among all cortical networks, SAL/PMN exhibits the weakest functional connectivity with both the action and the default networks, yet simultaneously displays the most pronounced spatial variability across cortical territories. At first glance, this weak coupling might suggest a marginal role. However, viewed through the lens of complex dynamical systems and the ancient philosophy of the Taijitu (Yin‐Yang diagram), this pattern elevates SAL/PMN to a uniquely pivotal position. It is analogous to the dynamic “fish-eye” (the two small dots) embedded within the Yin and Yang poles. In the Taijitu, the fish-eyes are not passive remnants but active, generative foci where the opposite force is seeded, enabling the perpetual transformation and rhythmic alternation between Yin and Yang.

Translating this metaphor to the brain, SAL/PMN sits precisely at the interstice between the externally oriented action network (Yang) and the internally oriented default network (Yin). Its weak average affinity to either pole does not denote irrelevance. Rather, it signifies a position of maximal dynamical freedom and sensitivity, poised to detect and amplify critical fluctuations that trigger state transitions. The high spatial variability of SAL/PMN connectivity is not a nuisance but a functional signature of its role as a “critical switching hub.” In dynamical systems theory, regions or modes that exhibit high variance and low mean coupling often operate near a critical point. Even small perturbations can produce large‐scale reconfigurations of the entire network state. This property endows SAL/PMN with the capacity to integrate interoceptive, exteroceptive, and cognitive signals, and to rapidly gate the dominance between action‐mode and default‐mode processing. This aligns with its known roles in salience detection, error monitoring, and autonomic arousal. Traditional TPC would likely obscure this unique variational signature. SPC clearly exposes it, underscores SPC’s superior ability to capture the dynamical essence of brain networks beyond static average strengths. SAL/PMN is not defined by how strongly it connects, but by how flexibly and variably it reconfigures its connectivity patterns across space and time, acting as the dynamic “eye” that orchestrates the rhythmic breathing between the brain’s two fundamental modes.

### Implications for AI and Future Directions

This finding carries profound implications for AI. Most current AI architectures rely on fixed, strongly weighted connections and optimize for stable, low‐variance representations. This makes them brittle when facing ambiguous or contradictory inputs that require rapid task‐switching. The SAL/PMN analogy suggests that a robust cognitive architecture may benefit from incorporating a dedicated, high‐entropy “switch” module. This module would maintain weak but extremely plastic connections to both the “action” and “contemplation” subsystems. Its internal dynamics would be characterized by high variability and sensitivity to context^10^. Such a module would not perform direct computation on inputs. Instead, it would continuously monitor the relative coherence and prediction errors of the two primary systems. It would trigger a cascade of reconfigurations whenever the environment demands a shift between external goal‐directed execution and internal simulation. This resonates with the concept of a “global neuronal workspace” and with the “contemplative system” driven by brain dark energy that we have previously posited^47,48^. By embracing rather than suppressing high spatial variability, future AI systems could achieve the kind of agile, context‐sensitive metastability that characterizes human cognition. The “fish-eye” of the system, though statistically weak, becomes the linchpin of adaptive intelligence, enabling seamless transitions, creative insight, and balanced energy allocation across competing computational regimes. The SPC framework, by faithfully revealing this hidden dynamical architecture, provides both neuroscientific validation and a computational blueprint for embedding genuine flexibility into the next generation of intelligent agents.

### Open Science and Concluding Remarks

In summary, the action–default anticorrelation, together with the SAL/PMN network, forms into a “three-body” system^61^ revealed by network SPC. This illuminates the energetic, computational, and evolutionary logic of brain organization. It offers a principled blueprint for designing next‐generation AI systems that are not only performant but also cognitively aligned with the human mind. The intriguing tripartite dynamics open several critical avenues for future investigation. In this framework, SAL/PMN acts as a highly variable “chaotic comet” or dynamic fish‐eye, weakly coupled to the other two networks yet exquisitely sensitive to perturbations. This positions the entire system at the edge of criticality for agile state transitions between external action and internal contemplation. Future work leveraging PET‐MRI across the entire lifespan is ideally suited to map the metabolic trajectories of this neural trio, from the ordered, periodic oscillations of development, through the optimally chaotic metastability of adulthood, to the potential breakdown or rigidification observed in aging and neurodegeneration. We will quantify how glucose and oxygen metabolic fluxes are dynamically reallocated among these three poles, testing the hypothesis that cognitive efficiency hinges on maintaining their delicate, chaotic equilibrium.

Moreover, this biological paradigm offers a transformative blueprint for next‐generation AI. Embedding a chaotic “modulator” module that mirrors SAL/PMN network’s high‐variability switching could endow deep learning systems with autonomous, energy‐efficient task‐switching and counterfactual imagination, pushing AI closer to the brain’s remarkable adaptability and parsimonious energy governance. Unraveling this cerebral three‐body interplay promises to unify systems neuroscience, metabolic imaging, and brain‐inspired computing under a single, coherent dynamical framework. It offers a more holistic and biologically grounded model of brain function and represents a paradigm shift in understanding how the brain orchestrates goal-directed behavior.

However, for such a promising framework to realize its full potential^62^, a strong commitment to open science is essential. Our work necessitates a concerted effort to publicly share: 1) standardized action network atlas resources, including high-resolution maps for different age groups and parcellation schemes; and 2) computational analysis code, including the full code used for generating the network maps, functional connectivity analyses, and statistical comparisons. This will allow researchers worldwide to confidently identify the action network in their own datasets, enabling independent validation and exploration across diverse populations and clinical conditions. Transparency in processing pipelines is critical for reproducibility and for allowing other teams to build upon this work efficiently. Embracing these open science principles will democratize access to the action network framework, empowering a broader range of scientists to test its predictions. This collaborative and transparent approach is the most effective way to transform the action network from a compelling theoretical model into a foundational tool for cognitive neuroscience and neurotherapy. Ultimately, open sharing of these resources will catalyze innovation, ensuring that the study of the brain’s action-mode benefits from the collective intelligence of the global scientific community^63^.

## Data availability statement

All codes and MIGP-derived data are deposited in the Chinese Color Nest Data Community (CCNDC) in the Science Data Bank (https://doi.org/10.57760/sciencedb.24589).

## Ethical statement

Data were provided [in part] by the Human Connectome Project (HCP), WU-Minn Consortium (Principal Investigators: David Van Essen and Kamil Ugurbil; 1U54MH091657) funded by the 16 NIH Institutes and Centers that support the NIH Blueprint for Neuroscience Research; and by the McDonnell Center for Systems Neuroscience at Washington University; and by the Chinese Human Connectome Project (CHCP: Principal Investigators: Jia-Hong Gao and Xi-Nia n Zuo; Z171100000117012 and Z181100001518003) funded by the Beijing Municipal Science and Technology Commission. All subjects provided written informed consent. All the protocols were approved by the Institutional Review Boards at Peking University (CHCP) and Washington University in Saint Louis (HCP). More details of the ethical statements can be found in their original publications^23,24^.

## Author contributions

X-XX: Conceptualization, Data curation, Formal analysis, Software, Visualization, Writing – original draft, review & editing. X-NZ: Formal analysis, Software, Visualization, Funding acquisition, Writing – review & editing.

## Funding

This work has been supported by the scientific and technological innovation 2030 - the major project of the Brain Science and Brain-Inspired Intelligence Technology (2021ZD0200500).

## Conflict of interest

There are no competing interests to declare.

## Notes

https://ccndc.scidb.cn/en

### Competing Interest Statement

The authors have declared no competing interest.

### Summary of Updates

A full revision of the original submission.

