## Supplementary Materials for "Spatial Connectivity Pattern of The Human Brain’s Action Network"

### Supplementary Materials and Methods

#### *Open Datasets & Preprocessing*

To replicate and validate our findings derived from CHCP[23], we use the open datasets from the Human Connectome Project (HCP)[24]. Four 15-min (LR and RL phase encodings for each day, respectively) rfMRI scans for all participants. All preprocessing steps of the rfMRI data were identical between CHCP and HCP with an updated version of the HCP pipeline (https://github.com/Washington-University/HCPpipelines/releases/tag/v4.0.0). After quality control, a total of 1003 healthy human young adults were included in the subsequent analysis and their preprocessed individual rfMRI data has been openly shared through the HCP website as described in the original publications[24].

#### *Group-level Temporal and Spatial Connectivity*

Each preprocessed individual dataset was temporally trimmed, and variance normalization was applied and submitted to the principal component analysis (PCA) at the group level (MIGP). The group PCA compressed the datasets with the weighted top 4500 spatial eigenvectors for further renormalization and reweight to form group-level timeseries (**.dtseries*). According to rfMRI protocols, these PCA-compressed MIGP time series capture intrinsic spontaneous brain activity at the group level (i.e. 54 minutes).

The MIGP-timeseries are then correlated to form the average dense TPC connectomes (**.dconn*) at the group level (91282x91282 entries). Specifically, $g_i$ denotes the $i$-th gray coordinate ($i = 1,2,\ldots,91282$), and $t_i = [ts(g_i, 1), ts(g_i, 2), \ldots, ts(g_i, 2500)]$ denotes standardized timeseries measured at the coordinate $g_i$. The TPC connectome's entry at $(g_i, g_j)$ is defined by the cosine similarity between $t_i$ and $t_j$ as the equation (1).

$$tpc(i,j) = \frac{1}{2}\ln\left(\frac{1 + t_i \cdot t_j}{1 - t_i \cdot t_j}\right) \qquad (1)$$

Here, $s_i = [tpc(i, 1), tpc(i, 2), \ldots, tpc(i, 91282)]$ denotes standardized full-brain TPC map seeded at the coordinate $g_i$. The spatial connectivity between $g_i$ and $g_j$ is defined by the cosine similarity between $s_i$ and $s_j$ as the equation (2).

$$spc(i,j) = \frac{1}{2}\ln\left(\frac{1 + s_i \cdot s_j}{1 - s_i \cdot s_j}\right) \qquad (2)$$

This derives the full-brain SPC connectome's entry at $(g_i, g_j)$ based on average dense TPC connectome at the group level. Specifically, we treat the full TPC profile of a given spatial location or element of volume as a snapshot of the traveling wave dynamics measured by the spontaneous slow oscillations (SSO) from rfMRI scans. We then standardized cosine angles between each pair of snapshots to produce the dense SPC connectome at the group level.

*Network Connectivity Mapping*

As illustrated in Figure S1, four seeds of the default network are selected for each hemisphere according to the DU15NET parcellation in the fsLR_32k HCP graycoordinate space including 91,282 elements (59412 for the cerebral cortex, 17853 for the cerebellum, 14017 for subcortical nuclei) and their spatial and label information are summarized in Table S1. Given a seed, its two-step neighbors' SPC maps were extracted from the group-level dense SPC connectomes. The median map of these SPC maps was calculated to quantify the seed network connectivity. All the eight seeds' network SPC maps were averaged as the default network SPC map. The default network TPC are also derived with the same approach for methodology comparison.

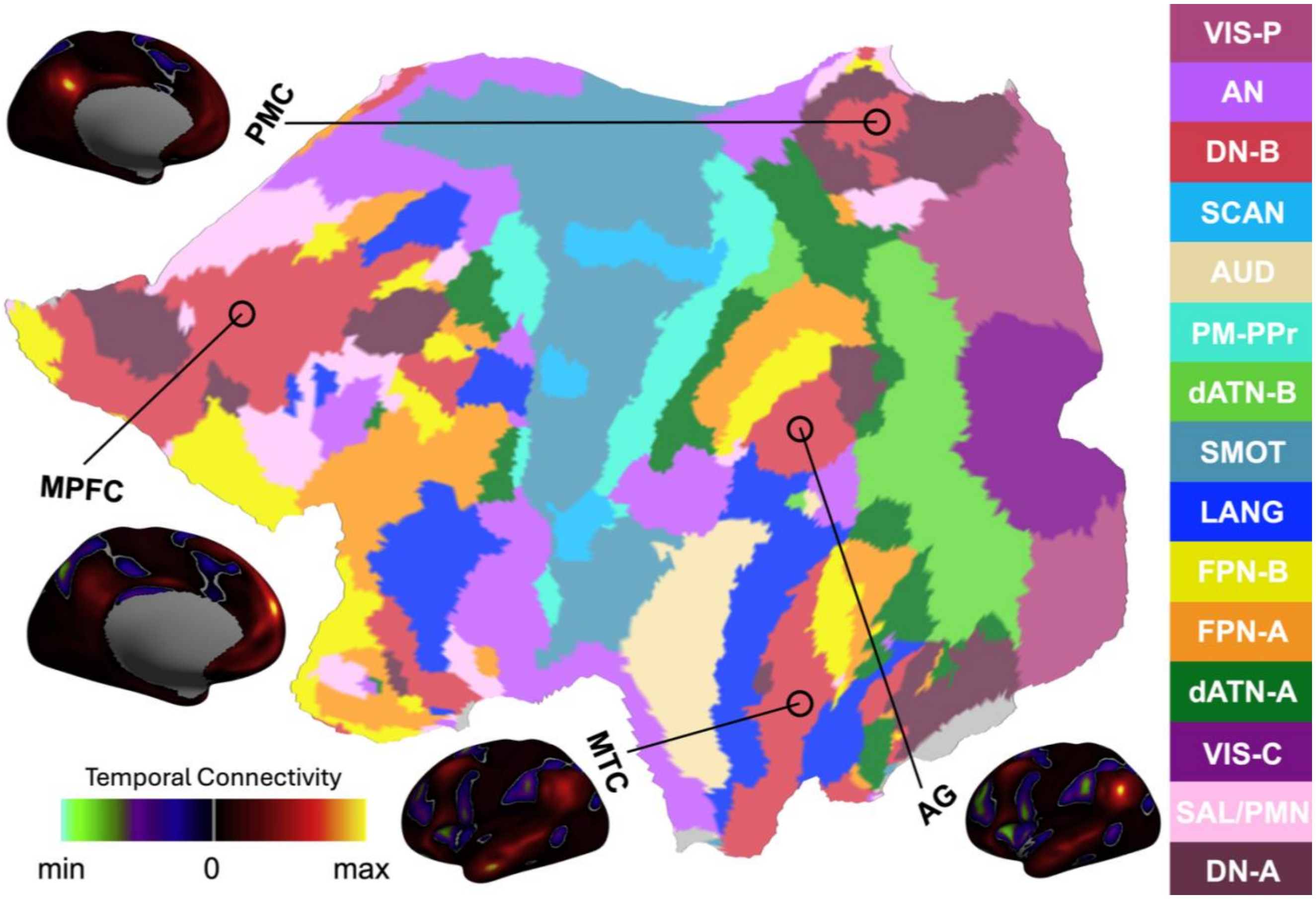


**Figure S1. Methodology of default network connectivity in the human brain**. Four seed regions are circled on the flat map of the left hemisphere: parietal medial cortex (PMC), medial prefrontal cortex

(MPFC), angular gyrus (AG) and medial temporal cortex (MTC). The original label of action network in DU15NET is cingulo-opercular (CG-OP) named by its anatomical structures network and has been updated by its functional correspondence ‘action network’ (AN). Meanwhile, the two somatomotor networks (SMOT-A and SMOT-B) are merged and then divided into the somato-cognitive action network (SCAN) and another remaining somatomotor network (SMOT). Other twelve networks are premotor-posterior parietal rostral (PM-PPr), salience/parietal memory network (SAL/PMN), dorsal attention-A (dATN-A), dorsal attention-B (dATN-B), frontoparietal network-A (FPN-A), frontoparietal network-B (FPN-B), default network-A (DN-A), default network-B (DN-B), language (LANG), visual central (VIS-C), visual peripheral (VIS-P), and auditory (AUD). The TPC maps of the four seed affiliated with DN-B areas are rendered onto the fsLR_32k cortical surfaces and visualized, respectively.

**Table S1. Information of the four seeds in the human default network**

| **full name** | **Posterior Medial Cortex** | **Medial Prefrontal Cortex** | **Angular Gyrus** | **Medial Temporal Cortex** |
|---|---|---|---|---|
| **abbreviation** | PMC | MPFC | AG | MTC |
| **XYZ (LH)** | -12.1, -51.5, 30.3 | -4.9, 58.1, 24.0 | -45.4, -59.3, 31.0 | -53.7, -9.5, -25.3 |
| **XYZ (RH)** | 13.0, -49.6, 31.4 | 7.0, 57.6, 21.4 | 50.3, -50.5, 29.9 | 48.1, -0.9, -30.8 |

### Supplementary Results

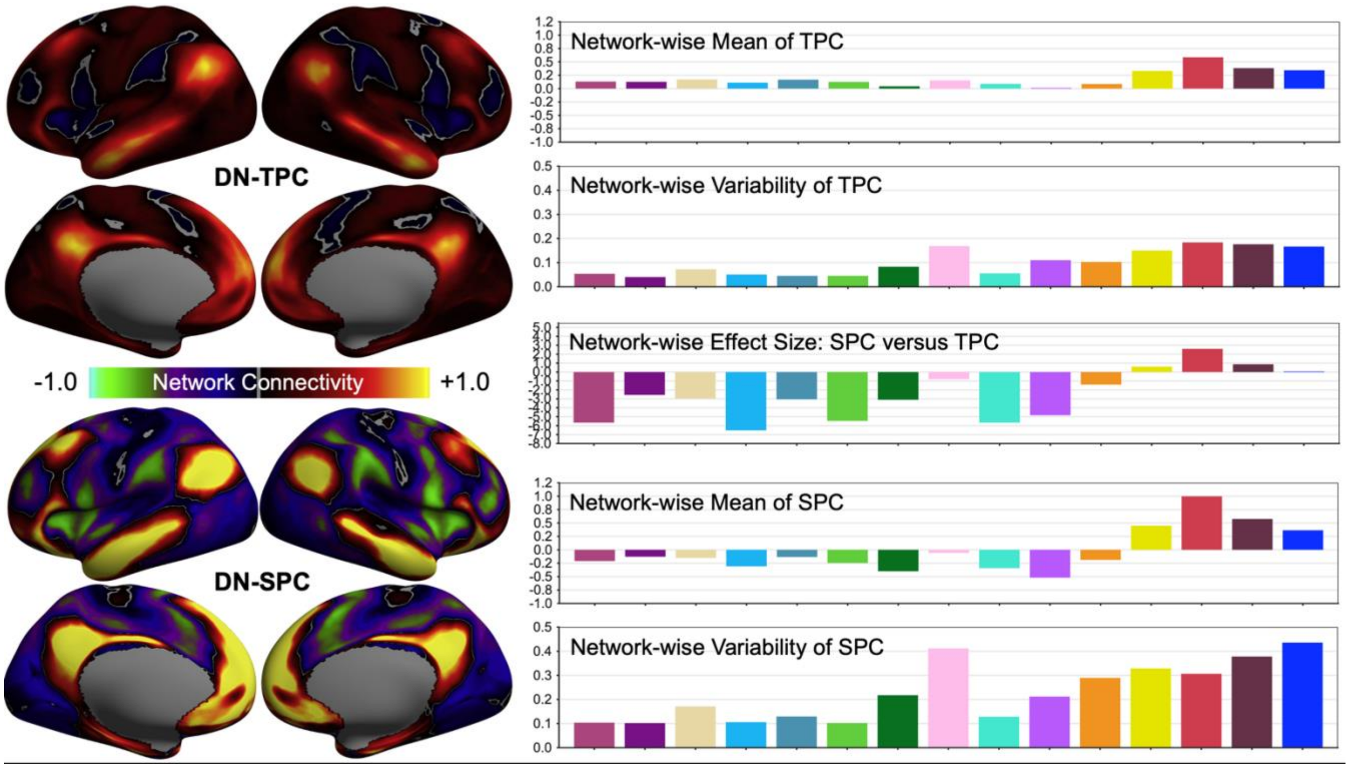


**Figure S2. Mapping default network connectivity in the human brain**. The network connectivity maps of action network (DN) are derived from the CHCP rfMRI data. Its temporal connectivity (DN-TPC) maps are rendered onto the standard cortical surfaces (top left). The network-wise mean or variability of TPC values are plotted as in the bar graph across the adjusted DU15NET networks (top right). Same surface maps and bar plots are presented for spatial connectivity of default network (DN-SPC) in bottom left and bottom right. To compare maps between TPC and SPC, we plot the network-wise effect size of SPC minus TPC in the middle right panel.

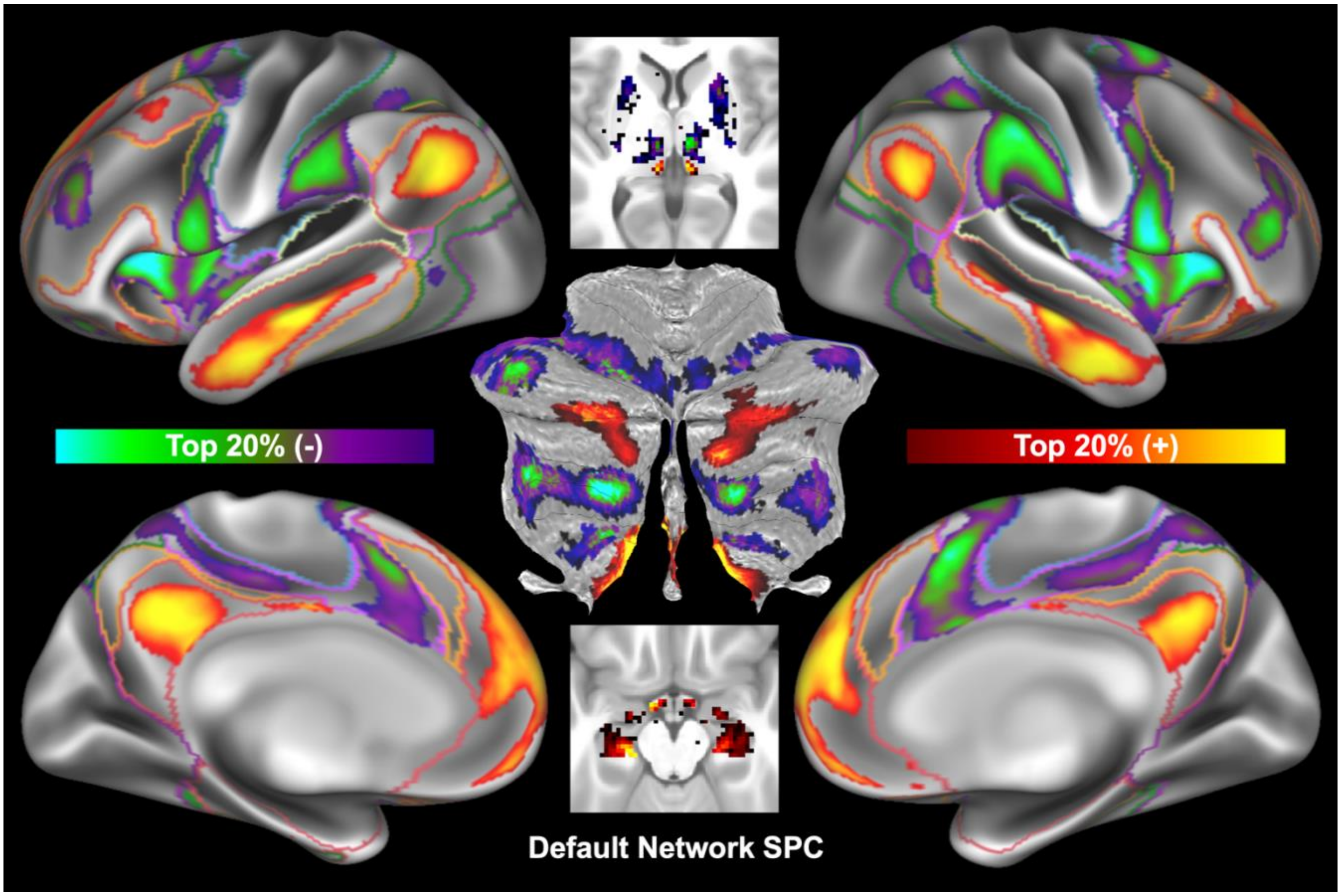


**Figure S3. Spatial connectivity pattern of default network in human brain**. The SPC map of default network was derived from the CHCP rfMRI data. Only the top 20% network SPC values were displayed for the three brain structures: cerebral cortex, subcortical nuclei and cerebellum. An integrative version of the DU15NET parcellation was used to outline boundaries of the seven large-scale networks derived by the CHCP team: visual (purple), somatomotor (blue), auditory (cream), dorsal attention (green), action (voliet), control (orange) and default (red). The default network SPC map was depicted in the two axial slices (z=4mm and z=-17mm) to demonstrate the high affinity of putamen and thalamus. This SPC map was also rendered on the flattened surface of the cerebellum.

### HCP Replications

Here, we presented the results derived from HCP dataset. Specifically, network connectivity maps of the human brain’s action network are shown in Figure S4. Traditional TPC showed only positive connectivity of the action network with the 15 cortical networks while varied across the networks. Specifically, sensorimotor, visual and auditory networks exhibited the strongest AN-TPC while default and control networks had the weakest AN-TPC. Interestingly, the most variable AN-TPC existed between action network and salience/parietal memory network. These observations are reproducible for AN-SPC. However, beyond the enhanced positive connectivity, AN-SPC produced obvious negative values (i.e., anticorrelation) in default network and control network. Such differences were further confirmed by the analysis of network-wise effect size on SPC-TPC as described in the equation (3). Similar findings are presented in Figure S5 for default network.

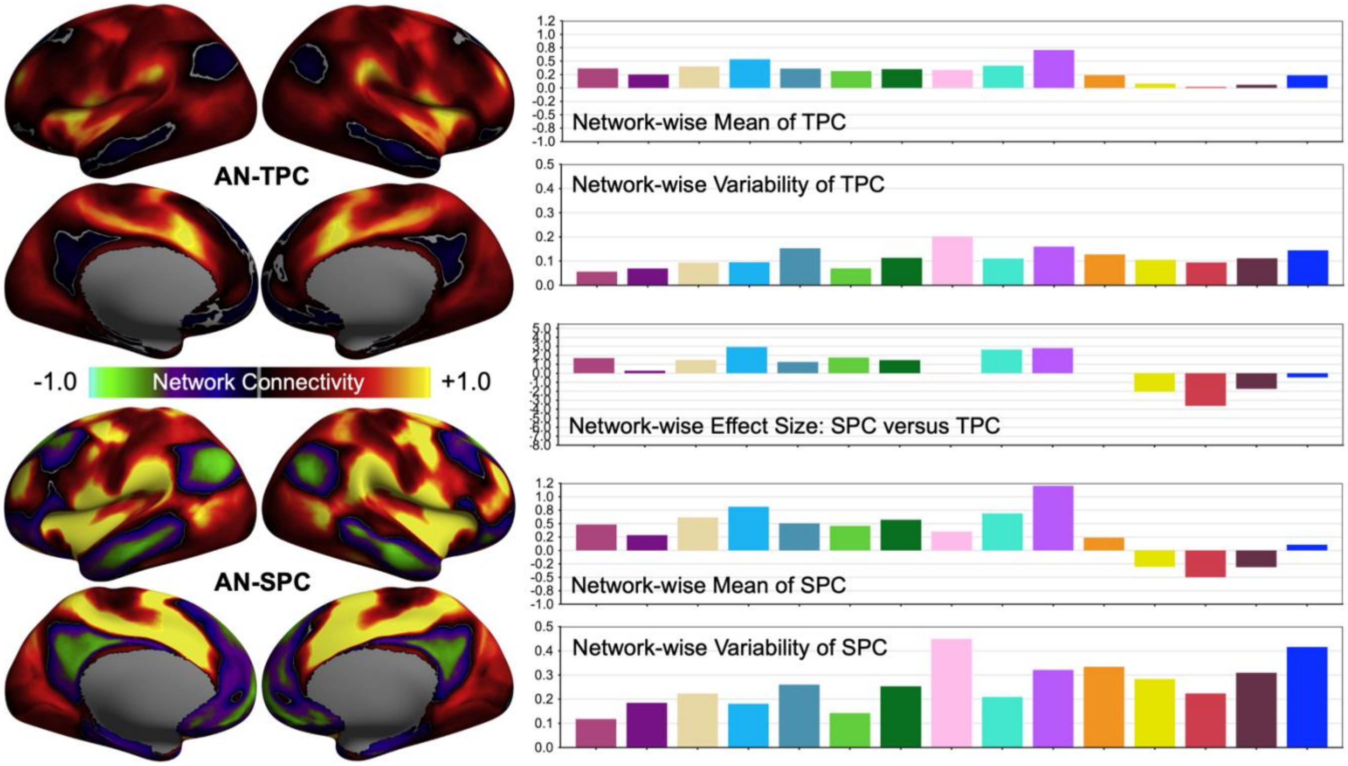


**Figure S4. Mapping action network connectivity in the human brain**. The network connectivity maps of action network (AN) are derived from the HCP rfMRI data. Its temporal connectivity (AN-TPC) maps are rendered onto the standard cortical surfaces (top left). The network-wise mean or variability of TPC values are plotted as in the bar graph across the adjusted DU15NET networks (top right): the same color annotation used as in Figure 1. Same surface maps and bar plots are presented for spatial connectivity of action network (AN-SPC) in bottom left and bottom right. To compare maps between TPC and SPC, we plot the network-wise effect size of SPC minus TPC in the middle right panel.

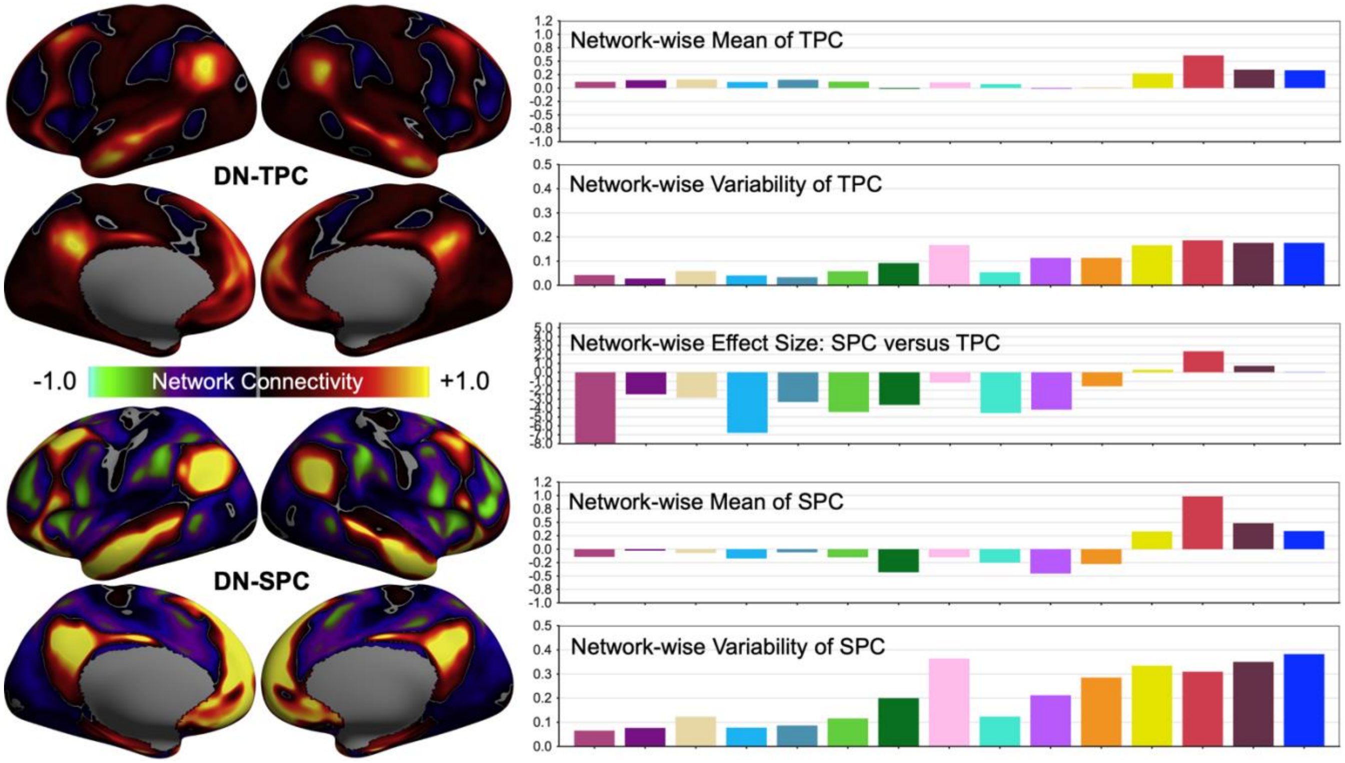


**Figure S5. Mapping default network connectivity in the human brain**. The network connectivity maps of default network (DN) are derived from the HCP rfMRI data. Its temporal connectivity (AN-TPC) maps are rendered onto the standard cortical surfaces (top left). The network-wise mean or variability of TPC values are plotted as in the bar graph across the adjusted DU15NET networks (top right): the same color annotation used as in Figure 1. Same surface maps and bar plots are presented for spatial connectivity of default network (DN-SPC) in bottom left and bottom right. To compare maps between TPC and SPC, we plot the network-wise effect size of SPC minus TPC in the middle right panel.

The upper 20% of the AN-SPC is visualized with positive or negative values to characterize synchronous or asynchronous functional communications of spatial-temporal dynamics in the cerebral cortex, cerebellum and subcortical nuclei, respectively (Figure S6). This method of mapping action network connectivity precisely delineates the cortical topology of the action network that outlines the boundaries (the violet color) of common networks, namely the cingulo-opercular or salience/ventral network[23,28,31]. We note that the negative connectivity of the action network is well distributed in the areas of the default network with red boundaries, echoing their 'Yin-Yang' communicative dynamics. This profile of integration of the action-default network was also mirrored in the cerebellum topology of the affinity of the action network, as well as in the affinity of the subcortical action network (mainly in the putamen and thalamus). The complete default network's SPC map is shown in Figure S7, and demonstrates a highly similar but mirror pattern of network spatial connectivity.

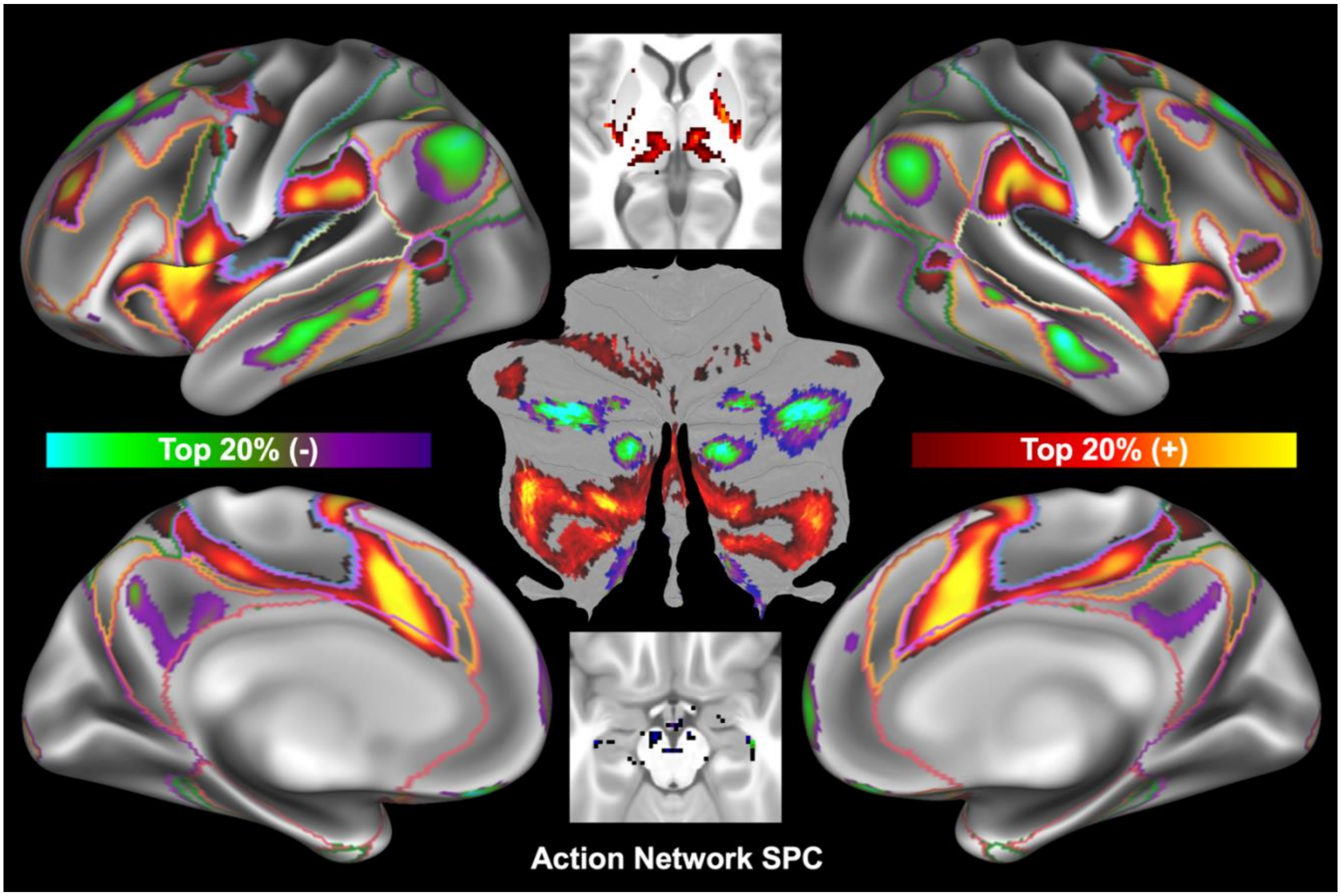


**Figure S6. Spatial connectivity pattern of action network in human brain**. The SPC map of action network was derived from the HCP rfMRI data. Only the top 20% network SPC values were displayed for the three brain structures: cerebral cortex, subcortical nuclei and cerebellum. An integrative version of the DU15NET parcellation was used to outline boundaries of the seven large-scale networks derived by the HCP team: visual (purple), somatomotor (blue), auditory (cream), dorsal attention (green), action (voliet), control (orange) and default (red). The action network SPC map was depicted in the two axial slices (z=4mm and z=-17mm) to demonstrate the high affinity of putamen and thalamus. This SPC map was also rendered on the flattened surface of the cerebellum.

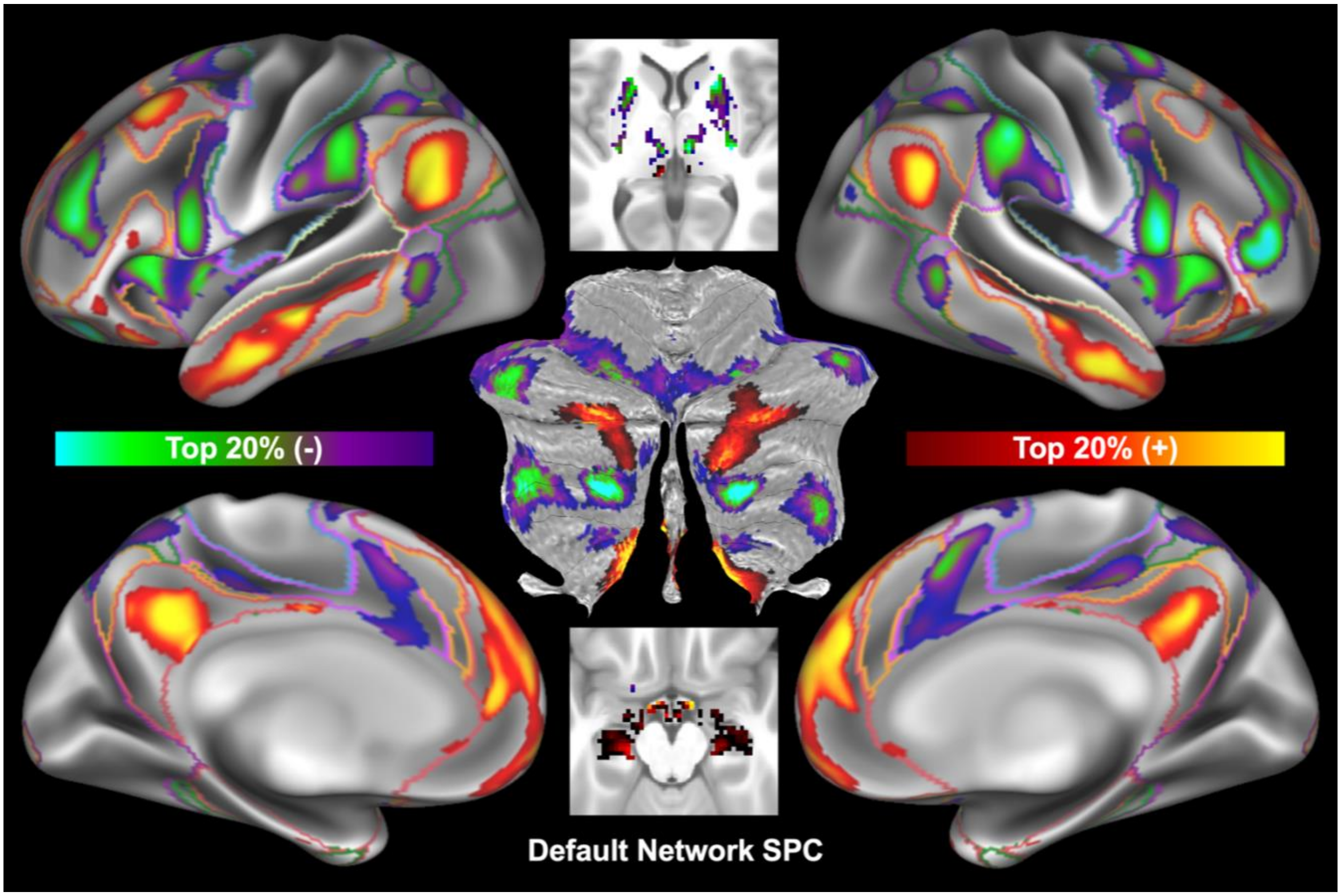


**Figure S7. Spatial connectivity pattern of default network in human brain**. The SPC map of default network was derived from the HCP rfMRI data. Only the top 20% network SPC values were displayed for the three brain structures: cerebral cortex, subcortical nuclei and cerebellum. An integrative version of the DU15NET parcellation was used to outline boundaries of the seven large-scale networks derived by the HCP team: visual (purple), somatomotor (blue), auditory (cream), dorsal attention (green), action (voliet), control (orange) and default (red). The default network SPC map was depicted in the two axial slices (z=4mm and z=-17mm) to demonstrate the high affinity of putamen and thalamus. This SPC map was also rendered on the flattened surface of the cerebellum.
